# Antibody-Dependent Heterotypic Syncytia Drive COVID-19 Inflammation and Disease Progression

**DOI:** 10.64898/2026.02.11.705426

**Authors:** Yin Yang, Chi Zhou, Tianyi Zhang, Weiqi Hu, Nicholas Magazine, Mariano Carossino, Nannan Jia, Teng Li, Yingdan Wang, Chen Wang, Jinghe Huang, Weishan Huang, Fan Wu

## Abstract

Severe COVID-19 is characterized by profound immune dysregulation and excessive inflammation. Aberrant myeloid responses drive hyperinflammation, yet the precise mechanisms remain elusive. Here, we demonstrate that certain spike antibodies promote the formation of heterotypic syncytia between monocytes/macrophages and virus-infected pneumocytes, leading to excessive inflammatory cytokine release. This antibody-dependent syncytium formation requires FcγRI (CD64) and ADAM10, as well as the assembly of the six-helix bundle by the spike S2 subunit. Notably, these syncytia exhibit a stronger proinflammatory signature compared to virus-infected epithelial cells. *In vivo*, sub-neutralizing antibody concentrations amplified inflammation and exacerbated disease in SARS-CoV-2-infected mice without increasing viral burden. Furthermore, scRNA-seq of postmortem lung tissues from COVID-19 patients implicated the monocyte/macrophage-derived syncytia as a key cellular source of inflammatory cytokines. Together, these findings define a novel mechanism of antibody-dependent enhancement of inflammation that helps explain the hyperinflammatory responses in severe COVID-19 and suggest new therapeutic opportunities targeting this pathway.

## INTRODUCTION

The COVID-19 pandemic caused by SARS-CoV-2 has resulted in over 779 million confirmed cases and more than 7.1 million deaths globally (1). While the majority of infected individuals exhibit mild disease or are asymptomatic, some patients develop severe pulmonary damage and acute respiratory distress syndrome (ARDS), which can progress to multi-organ failure and death. The prevailing view is that the major cause of severe COVID-19 pathogenesis is the excessive inflammatory response resulting from immune dysregulation, rather than direct damage caused by viral infection (2). However, the mechanisms underlying this hyperinflammation remain poorly understood.

Postmortem examinations of lungs have revealed frequent vascular injury, diffuse alveolar damage, moderate fibrosis, and extensive infiltration of inflammatory monocytes and macrophages. Notably, pneumocyte syncytia harboring SARS-CoV-2 RNA were commonly observed in these patients (3, 4) and are thought to contribute to disease severity by promoting cell-to-cell viral spread (5). Consistent with the histologic findings, scRNA-seq analyses of autopsy and BALF samples from patients with severe COVID-19 confirmed extensive infiltration of monocytes and macrophages and showed that these cells contained even more SARS-CoV-2 RNA than epithelial cells, along with high expression of pro-inflammatory cytokines (6, 7). Importantly, we and others have reported that SARS-CoV-2 neutralizing antibody titers correlate positively with COVID-19 severity (8), and that the pathogenic effects of antibodies are modulated by their Fc afucosylation status, which influences antibody interactions with Fcγ receptors (FcγRs) (9–11). Although human monocytes and macrophages are typically not susceptible to SARS-CoV-2 due to the lack ACE2 expression, anti-spike antibodies can mediate infection of these cells through FcγR binding (12–14). Thus, there is broad consensus that aberrantly activated monocytes and macrophages produce excessive amounts of inflammatory cytokines and are key drivers of the “cytokine storm” in severe COVID-19. However, the mechanisms of infection and activation of these cells remain unclear. More critically, the precise role of anti-spike antibodies in monocyte/macrophage-mediated inflammation has yet to be elucidated, particularly within the microenvironment where myeloid cells interface with pneumocytes.

In this study, we describe a previously unrecognized role of SARS-CoV-2 antibodies in shaping disease outcomes. We found that certain spike-specific antibodies promote syncytium formation between monocyte/macrophage and virus-infected pneumocyte, leading to robust release of inflammatory cytokines. Using both *in vitro* systems and animal models, we demonstrate that sub-neutralizing antibody concentrations can amplify inflammation and worsen disease severity without increasing viral burden. This antibody-mediated heterotypic cellular fusion requires FcγRI (CD64) and ADAM10 expression on monocytes, as well as membrane fusion mediated by the spike S2 subunit. These results define a previously unrecognized mechanism of antibody-dependent enhancement of inflammation that helps explain the excessive immune activation observed in severe COVID-19 and suggest new therapeutic opportunities targeting this process.

## RESULTS

### Spike NAb exacerbates disease severity in FcγRI-humanized mice infected with SARS-CoV-2

We previously screened 245 donors who had recovered from COVID-19 and isolated a broadly neutralizing antibody named GW01 from a donor whose plasma showed an extremely high titer of SARS-CoV-2 NAb (8, 15). GW01 targets a highly conserved receptor-binding domain (RBD) of spike and provides protection in mice during SARS-CoV-2 infection (15). Since the functions of antibodies depend not only on antigen binding but also, and more critically, on engagement with distinct types of Fc receptors (FcRs) of diverse immune cells (16, 17), we evaluated the protection of GW01 in FcγRI-humanized mice infected with mouse-adapted SARS-CoV-2 strain MA10 (18) (Figure 1A). Consistent with previous reports, GW01 efficiently inhibited viral replication and significantly reduced viral burden in infected mice (Figure 1C). However, administration of GW01 at an intermediate dose unexpectedly exacerbated weight loss relative to MA10 infection only group across time points (4 µg, *P* = 0.012) (Figure 1B). At 5 and 6 d.p.i., both the 4 µg and 40 µg GW01 treatment groups showed significantly greater weight loss than the infection-alone group (Figure 1B). Correspondingly, mice treated with 4 μg of GW01 showed significantly increased levels of IL-6, TNF, and CXCL12 expression compared with the MA10-infection-only group (Figure 1D). Lung tissues from infected mice treated with 4 μg of GW01 also showed a trend of higher pathologic scores in the parabronchial compartment (Figure 1E). These data suggest that at certain doses, GW01 induces inflammation and exacerbates disease severity *in vivo* despite efficiently neutralizing SARS-CoV-2.

**Figure 1.**
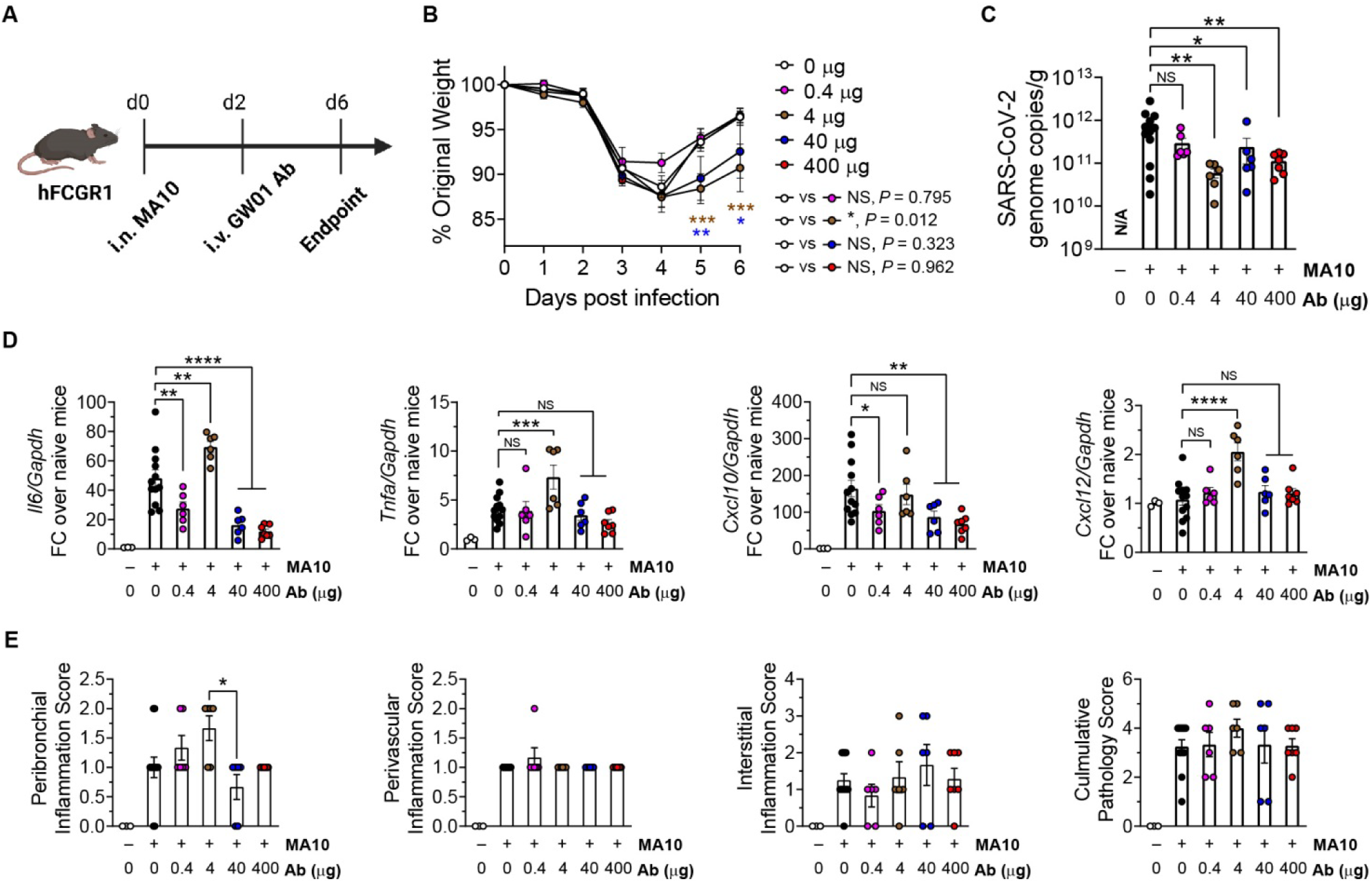
Spike NAb exacerbates disease severity in FcγRI-humanized mice during SARS-CoV-2 infection. (**A**) Schematic of the mouse infection and dosing regimen. (**B**) Body weight changes over time by GW01 dose. (**C**) Quantification of SARS-CoV-2 genome copies in lungs at endpoint. (**D**) Cytokine mRNA levels in lung homogenates by qPCR. (**E**) Pathologic scores of lung sections. Data are presented as mean ± SEM, n=12 for MA10 infection only group, n=6 for 0.4µg, 4µg and 40µg groups, n=7 for 400µg group. NS=not significant, *P<0.05, **P<0.01, ***P<0.001, ****P<0.0001, by one-way ANOVA.

### Spike NAb mediates syncytium formation between monocyte/macrophage and pneumocyte

We next investigated how GW01 causes excessive inflammation in SARS-CoV-2-infected FcγRI-humanized mice. During SARS-CoV-2 infection, myeloid cells, especially monocytes and macrophages, are critical sources of inflammatory cytokine (19). In addition, spike antibodies can mediate SARS-CoV-2 infection of monocytes and macrophages (12, 13). We generated a spike-expressing human A549 cell line (A549-spike) and co-cultured them with human monocytic THP-1 cells, with or without addition of GW01. We observed that the presence of GW01 induced multinucleated syncytia in an overnight co-culture (Figure 2A). To further visualize the syncytium formation process, we generated an mCherry-expressing THP-1 cell line and labeled A549-spike cells with CFSE, then cultured the two populations in the presence or absence of GW01. Live cell imaging (Figure 2B, Supplementary Figure 1, and Movie S2) revealed that GW01 induced heterotypic fusion between THP-1 cells (mCherry; Red) and A549-spike cells (CFSE; Green) within 3 hours following the addition of GW01. No fusion was observed in the absence of GW01 (Figure 2B, Supplementary Figure 1, and Movie S1). At 1 µg/mL, GW01-induced syncytia (double positive for eFluor450 and CFSE) constituted 10.1 % of the total cell population (Figure 2C). Notably, introducing LALA mutations into the GW01 heavy chain to disrupt Fc binding abrogated fusion between THP-1 and A549-spike cells, indicating that this process requires Fc receptor engagement on THP-1 cells (Figure 2C). We next asked whether antibodies could induce syncytium formation in primary human cells. Primary monocytes formed syncytia at levels comparable to THP-1 cells (Figure 2, D and E). We further differentiated monocytes with GM-CSF or M-CSF to generate monocyte-derived macrophages (MDMs), named G-MDMs and M-MDMs, respectively. G-MDMs formed syncytia, whereas M-MDMs showed markedly reduced syncytium formation (Figure 2, F and G). Phenotypically, G-MDMs resemble human alveolar macrophages, whereas M-MDMs are more similar to peritoneal macrophages (20), suggesting that the lung microenvironment might be specifically associated with antibody-mediated syncytium formation between monocyte/macrophage and infected pneumocyte.

**Figure 2.**
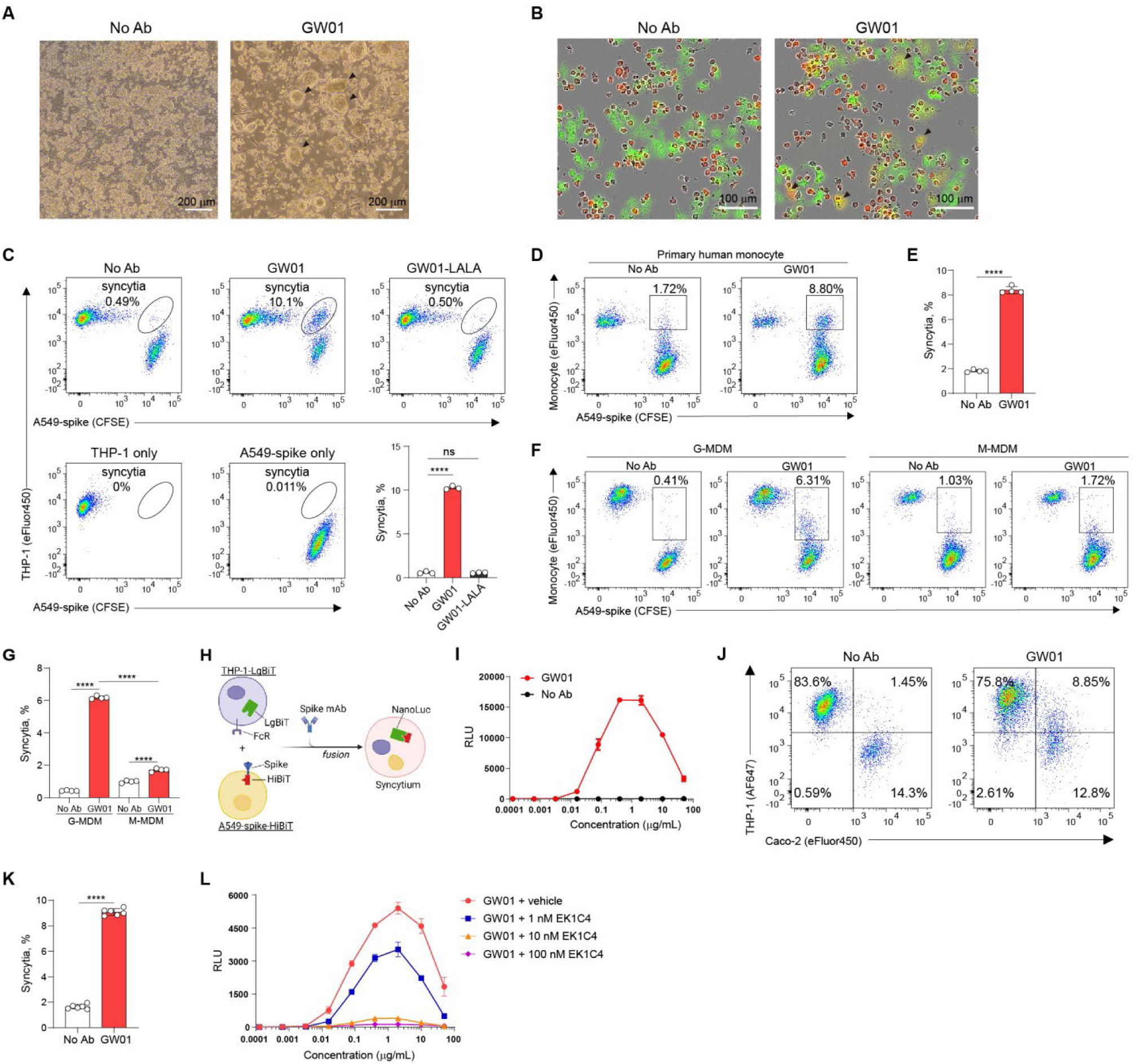
Spike NAb induces heterotypic syncytium formation between monocyte/macrophage and pneumocyte. (**A)** Bright-field images of THP-1 and A549-spike after 24-hour co-culture, with or without GW01. Scale bar: 200 μm. (**B**) Representative fluorescent micrographs of syncytia (yellow) between THP-1-mCherry (red) and A549-spike (green, labeled with CFSE) at 3 hours post co-culture. Scale bar: 100 μm. (**C**) Representative flow cytometry dot plots of GW01-induced syncytia (double positive population) between THP-1 and A549-spike under the indicated conditions. Quantification (bottom right) of syncytia (mean ± SEM, n=3, \*\*\*\**P*<0.0001). (**D**) Representative flow cytometry dot plots of GW01-induced syncytia between primary human moncytes and A549-spike. (**E**) Quantification of syncytia from (D) (mean ± SEM, n=4, \*\*\*\**P*<0.0001). (**F**) Representative FACS dot plots of GW01-induced syncytia between human monocyte-derived macrophages (MDMs) and A549-spike. MDMs were differentiated with either GM-CSF (G-MDM) or M-CSF (M-MDM). (**G**) Quantification of syncytia from (F) (mean ± SEM, n=4, ****P<0.0001). (**H**) Experimental design of NanoLuc assay to test cell-cell fusion. (**I**) NanoLuc assay analysis of GW01 concentration-dependent effects on syncytium formation. **(J**) Syncytium formation during SARS-CoV-2 infection. (**K**) Quantification of syncytia from (J) (mean ± SEM, n=6, \*\*\*\**P*<0.0001). (**L**) Analysis of the effect of EK1C4 on syncytium formation.

To further determine how antibody concentration affects syncytium formation, we established a precise, high-throughput assay of heterotypic cell-cell fusion using HiBiT protein tagging (Figure 2H). Briefly, the HiBiT epitope (VSGWRLFKKIS) was fused to the C terminus of spike in A549-spike-HiBiT cells, and the complementary LgBiT subunit was stably expressed in THP-1-LgBiT cells. When heterotypic fusion occurs between A549-spike-HiBiT and THP-1-LgBiT cells, HiBiT and LgBiT assemble into NanoLuc, a functional luciferase that can be used for quantification by luminescence. Using this system, we observed that GW01-mediated fusion between A549-spike and THP-1 cells depended on the antibody concentration (Figure 2I): significant fusion was detected between 10 ng/mL and 100 µg/mL, peaking at 1 µg/mL, with no fusion above 100 µg/mL or below 1 ng/mL, consistent with the dose-dependent disease-exacerbating effect of GW01 *in vivo* (Figure 1A-E).

We next evaluated additional anti-spike monoclonal antibodies (Supplementary Figure 2). Among 21 RBD-specific antibodies tested, 6 (9L6, REGN10933, COVA2-15, 6M6, BD55-5514, and CAB-A17) induced detectable syncytium formation between THP-1 and A549-spike cells. Remarkably, all active antibodies exhibited a similar bell-shaped concentration dependence (Supplementary Figure 2), suggesting a conserved mechanism of action.

To test whether antibodies could mediate fusion of monocytes with cells infected by live viruses, we infected Caco-2-3a-E cells with SARS-CoV-2-ΔORF3a-E mNG virus (ΔORF3a-E mNG) (21) and co-cultured them with THP-1 cells in the presence of antibodies. The ΔORF3a-E mNG virus was engineered by deleting the ORF3a-E genes and inserting the mNeonGreen (mNG) reporter, such that the infected cells express green fluorescence. ΔORF3a-E mNG replication is supported in the complementing Caco-2-3a-E cell line. Here, Caco-2-3a-E cells were labeled with eFluor450, and THP-1 cells were stained with anti-CD64-Alexa Fluor 647 (AF647). Syncytia were quantified by flow cytometry as dual-positive events (eFluor450^+^ AF647^+^). In this live virus system, the spike-specific antibody GW01 induced heterotypic fusion between THP-1 cells and SARS-CoV-2-infected Caco-2 cells (Figure 2, J and K).

During spike-mediated fusion of viral and cellular membranes, binding of the spike RBD to ACE2 exposes the S2′ site, which is then cleaved by TMPRSS2 (22). Following cleavage, the fusion peptide inserts into the target membrane, and interactions between heptad repeat 1 (HR1) and HR2 appose the membranes to promote fusion (23). The peptide EK1C4 binds to HR1 and specifically inhibits spike-induced fusion (24). Using our HiBiT-based syncytium assay, EK1C4 inhibited antibody-mediated fusion in a concentration-dependent manner (Figure 2L). EK1C4 at 1 nM reduced fusion by ∼30%, whereas 100 nM completely abrogated syncytium formation. These findings indicate that antibody-mediated syncytium formation likewise requires the canonical spike-driven membrane-fusion machinery.

### Heterotypic syncytia exhibit a hyperinflammatory profile

To investigate how SARS-CoV-2 spike antibody-induced syncytium formation shapes inflammatory responses, we employed an *in vitro* co-culture system in which Caco-2-3a-E cells were infected with ΔORF3a-E mNG for 24 h, followed by the addition of GW01 and THP-1 cells (Figure 3A). At 24h after co-culture, we sorted fused syncytial cells, unfused THP-1 cells, and Caco-2-3a-E cells by FACS and performed RNA-seq. Pathway enrichment analysis revealed a marked upregulation of inflammatory pathways in both fused syncytial cells and THP-1 cells from the GW01-treated condition (Figure 3B), whereas ΔORF3a-E mNG-infected Caco-2-3a-E cells showed only minimal enrichment of these pathways (Figure 3B). In the absence of GW01, THP-1 cells displayed moderate enrichment of inflammatory pathways, likely reflecting direct stimulation of PRRs on the surface of THP-1 cells by SARS-CoV-2, such as TLR-2 and TLR-1 (25, 26) and paracrine cytokine signaling from infected Caco-2 cells. Together, these results indicate that antibody-mediated monocyte-derived syncytia display a robust pro-inflammatory signature, highlighting their potential role in driving inflammatory responses during infection.

**Figure 3.**
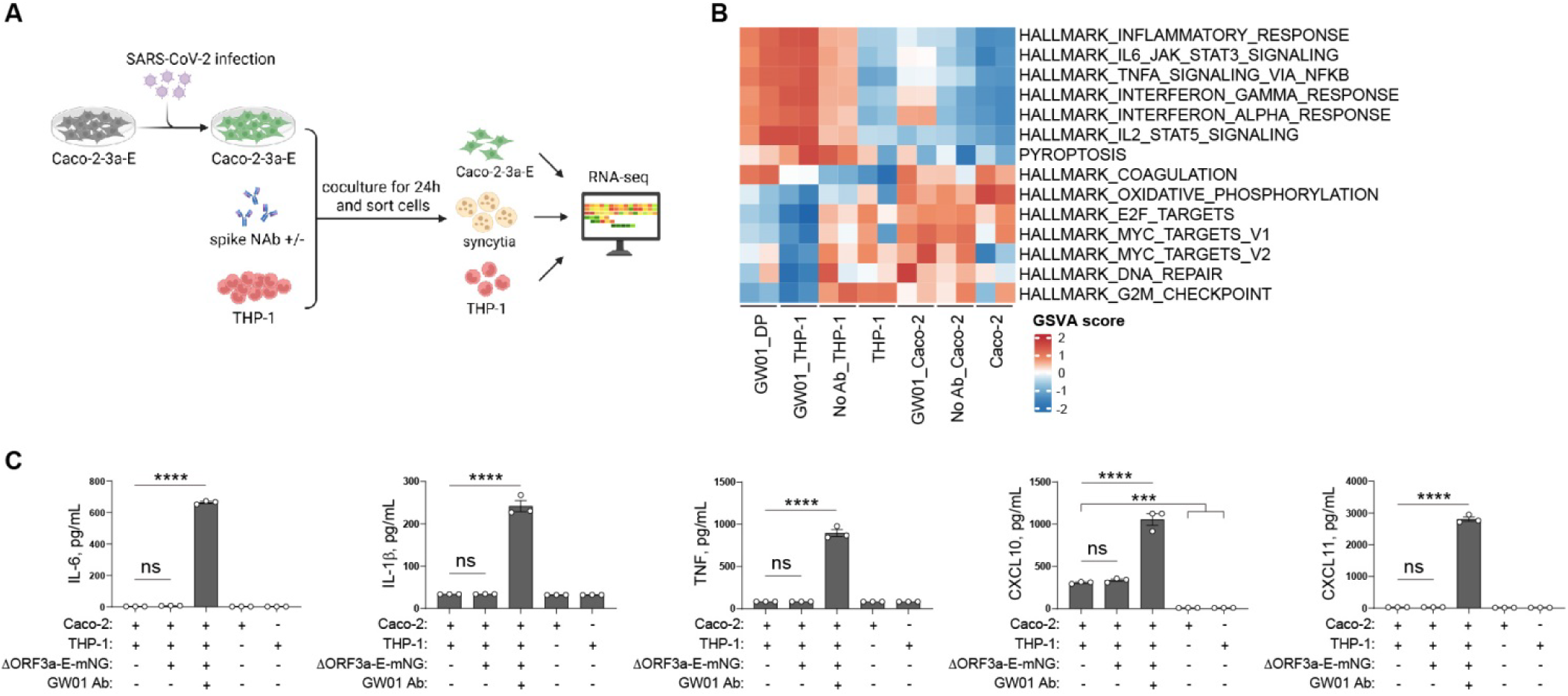
Antibody-induced syncytia during SARS-CoV-2 infection are highly pro-inflammatory. (**A**) Schematic of the co-culture assay used to characterize syncytial function. (**B**) Heatmap of pathway enrichment across the indicated cell populations. (**C**) Cytokine concentration in 24-h co-culture supernatants from (A), measured by ELISA. Data are presented as mean ± SEM, n=3; ns, not significant, ***P<0.001, ****P<0.0001, by one-way ANOVA.

Using the same co-culture assay, we quantified representative inflammatory cytokines and chemokines in supernatants. When ΔORF3a-E mNG-infected Caco-2-3a-E cells were co-cultured with THP-1 cells, GW01 elicited substantial production of IL-6, IL-1β, TNF, CXCL10, and CXCL11 (Figure 3C), whereas these inflammatory mediators were significantly reduced without GW01, mirroring GW01-enhanced inflammation in the humanized mouse infection model (Figure 1D). These findings indicate that spike antibody-induced monocyte-epithelial cell syncytia are critical contributors to the observed inflammatory responses.

### FcγRI and ADAM10 are critical for antibody-mediated heterotypic syncytium formation

Monocytes lack ACE2 or TMPRSS2, which are required for SARS-CoV-2 entry. In an attempt to identify host factors responsible for antibody-mediated fusion of monocytes with infected pneumocytes, we performed a CRISPR-Cas9 whole-genome knockout screen (Figure 4A). THP-1 cells stably expressing Cas9 were generated by lentiviral transduction and then transduced with a lentiviral sgRNA library covering the human genome. The resulting THP-1-Cas9 cells were co-cultured with antibody-coated A549-spike cells. Syncytia and non-fused THP-1-Cas9 cells were sorted by flow cytometry, and sgRNAs were sequenced. We found that sgRNAs targeting *FCER1G* and *ADAM10* were significantly enriched in non-fused cells, suggesting that both molecules are critical for syncytium formation (Figure 4B).

**Figure 4.**
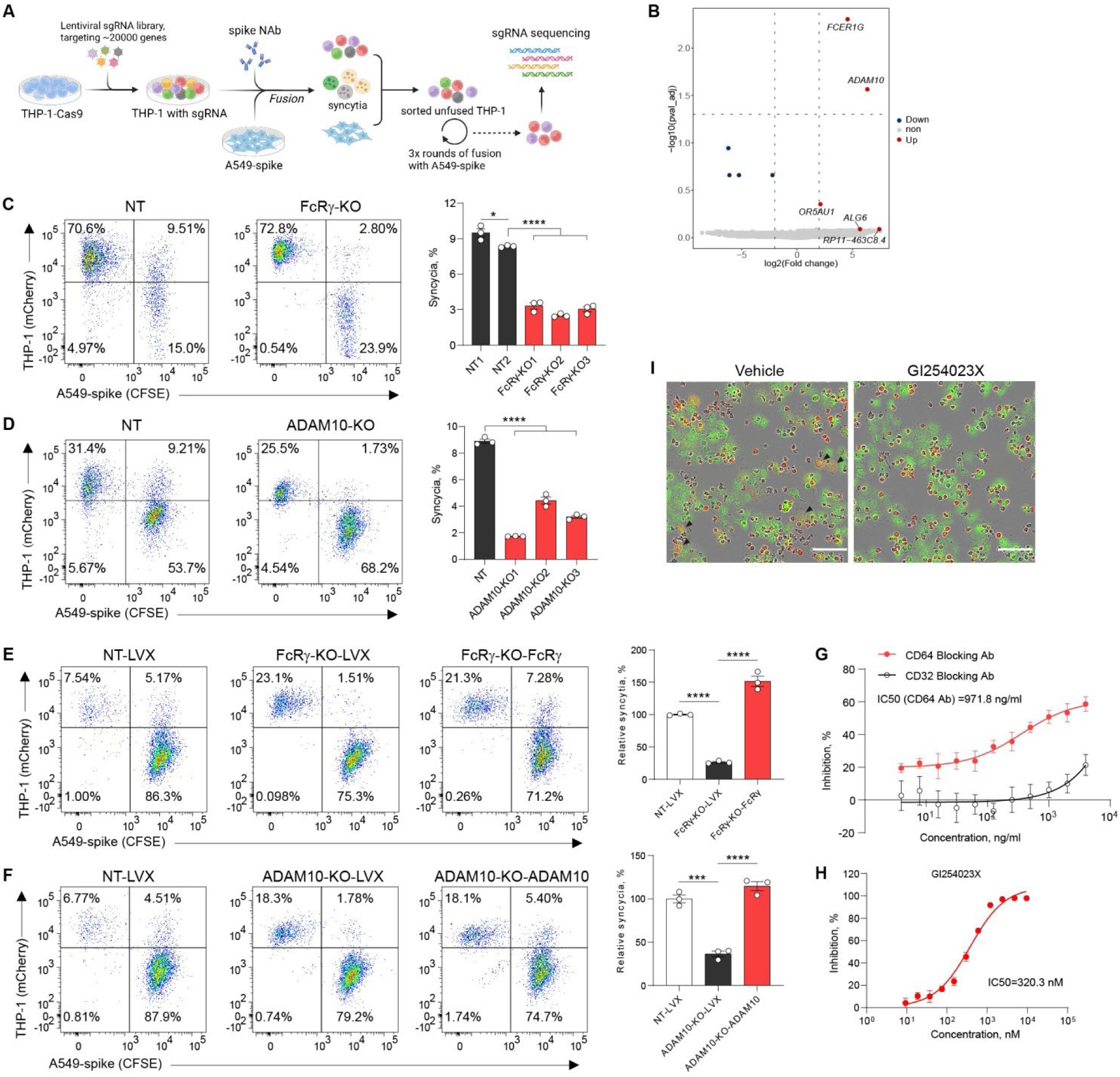
Monocyte FcRγ and ADAM10 are critical for antibody-dependent syncytium formation. (**A**) Schematic of the CRISPR-Cas9 screen. (**B**) Volcano plot of genes from the screen, ranked by fold change and adjusted *P* value. (**C**) Representative flow cytometry dot plots of GW01-induced syncytia between A549-spike and FcRγ-knockout (FcRγ-KO) THP-1 clones versus non-targeting control (NT), and quantification of syncytia (right inset) (mean ± SEM, n=3, \**P*<0.05, \*\*\*\**P*<0.0001). (**D**) Representative plots of A549-spike fusion with ADAM10-knockout (ADAM10-KO) THP-1 clones versus control and quantification of syncytia. (**E**) Rescue of FcRγ: representative plots for FcRγ-KO-LVX (FcRγ-knockout cells reconstituted with empty lentiviral vector), FcRγ-KO-FcRγ (FcRγ-knockout cells reconstituted with FcRγ cDNA), and NT-LVX (non-targeting control with empty lentiviral vector), with quantification of syncytia. (**F**) Rescue of ADAM10: representative plots for ADAM10-KO-LVX (ADAM10-knockout cells reconstituted with empty lentiviral vector), ADAM10-KO-ADAM10 (ADAM10-knockout cells reconstituted with ADAM10 cDNA), and NT-LVX, along with quantification of syncytia. (**G**) Effect of CD64 or CD32 blocking antibodies on GW01-dependent syncytium formation measured by NanoLuc assay. (**H**) Effect of the ADAM10 inhibitor GI254023X on GW01-dependent syncytium formation. (**I**) Representative fluorescence images of syncytia (yellow merge) between THP-1-mCherry (red) and A549-spike (green; CFSE) at 3 hours post co-culture, with or without GI254023X. Scale bar: 100 μm. Data are presented as mean ± SEM, n=3, \**P*<0.05, \*\*\**P*<0.001, \*\*\*\**P*<0.0001 by one-way ANOVA.

*FCER1G* encodes the common Fc receptor γ chain (FcRγ, also called FcεR1γ; hereinafter referred to as FcRγ), a signaling subunit of Fcγ receptor I (CD64) and Fcγ receptor III (CD16). FcRγ is essential for CD64 expression and function (27). Flow cytometry confirmed that FcRγ knockout abrogated CD64 surface expression on THP-1 cells (Supplementary Figure 3A). Consistent with a requirement for Fcγ receptors, GW01 bearing LALA mutations in the Fc region (GW01-LALA), which disrupt Fc-receptor binding, failed to induce syncytia (Figure 2C). To validate this finding, we generated three independent FcRγ-knockout THP-1 cell lines, all of which showed significantly impaired syncytium formation (Figure 4C), while reintroduction of FcRγ restored function (Figure 4E and Supplementary Figure 3A). Because THP-1 cells express two Fcγ receptors, CD64 and CD32, and only CD64 is associated with the FcRγ, we hypothesized that CD64, but not CD32, mediates antibody-induced syncytia. Indeed, anti-CD64 antibodies inhibited syncytium formation in a concentration-dependent manner (IC₅₀ = 971.8 ng/mL), whereas anti-CD32 had no effect (Figure 4G). These results establish CD64 as the critical FcγR for spike antibody–induced syncytium formation.

ADAM10 is a metalloproteinase best known for its sheddase activity on membrane proteins and its role in activating Notch signaling (28), but its role in antibody-mediated fusion has not been defined. Three independent ADAM10-knockout THP-1 cell lines showed impaired syncytium formation (Figure 4D), and reintroduction of ADAM10 fully restored syncytium formation (Figure 4F and Supplementary Figure 3B). Pharmacological inhibition with a selective ADAM10 inhibitor, GI254023X, confirmed these results, showing dose-dependent inhibition of syncytia with an IC₅₀ of 320.3 nM (Figure 4, H and I, Movie S3).

Together, these data demonstrate that both FcγRI (CD64, via FcRγ) and ADAM10 are required for antibody-induced heterotypic syncytium formation.

### ADAM10 is dispensable for FcγRI/antibody-mediated SARS-CoV-2 infection of monocytes via endocytosis

Recent studies have shown that spike antibodies can mediate SARS-CoV-2 infection of human monocytes via Fc receptors (12, 13), but the underlying mechanism remains unclear. Having demonstrated that FcγRI and ADAM10 are required for antibody-mediated cell–cell fusion, we next asked whether ADAM10 is also involved in the FcγRI/antibody-mediated viral infection. We used the SARS-CoV-2-ΔORF3a-E mNG virus, which can undergo only a single round of replication in host cells lacking ORF3a (21), to evaluate antibody-mediated infection in THP-1 cells. At a suboptimal antibody concentration of 0.2 ug/mL, the spike antibody GW01 mediated infection of THP-1 cells at a frequency of 7.8% (Figure 5A). Disrupting the GW01–FcγRI interaction with the LALA mutant (GW01-LALA) abolished the infection (Figure 5A). Consistent results were obtained with FcRγ-knockout THP-1 cells (Figure 5B), indicating that FcγRI is required for antibody-mediated infection of THP-1 cells. Unexpectedly, the ADAM10 inhibitor GI254023X had no effect at any dose tested (Figure 5A), and ADAM10-KO THP-1 cells showed similar infection levels to controls (Figure 5B). Thus, in contrast to antibody-mediated heterotypic cell-cell fusion, ADAM10 is not required for the antibody-mediated viral infection of monocytes.

**Figure 5.**
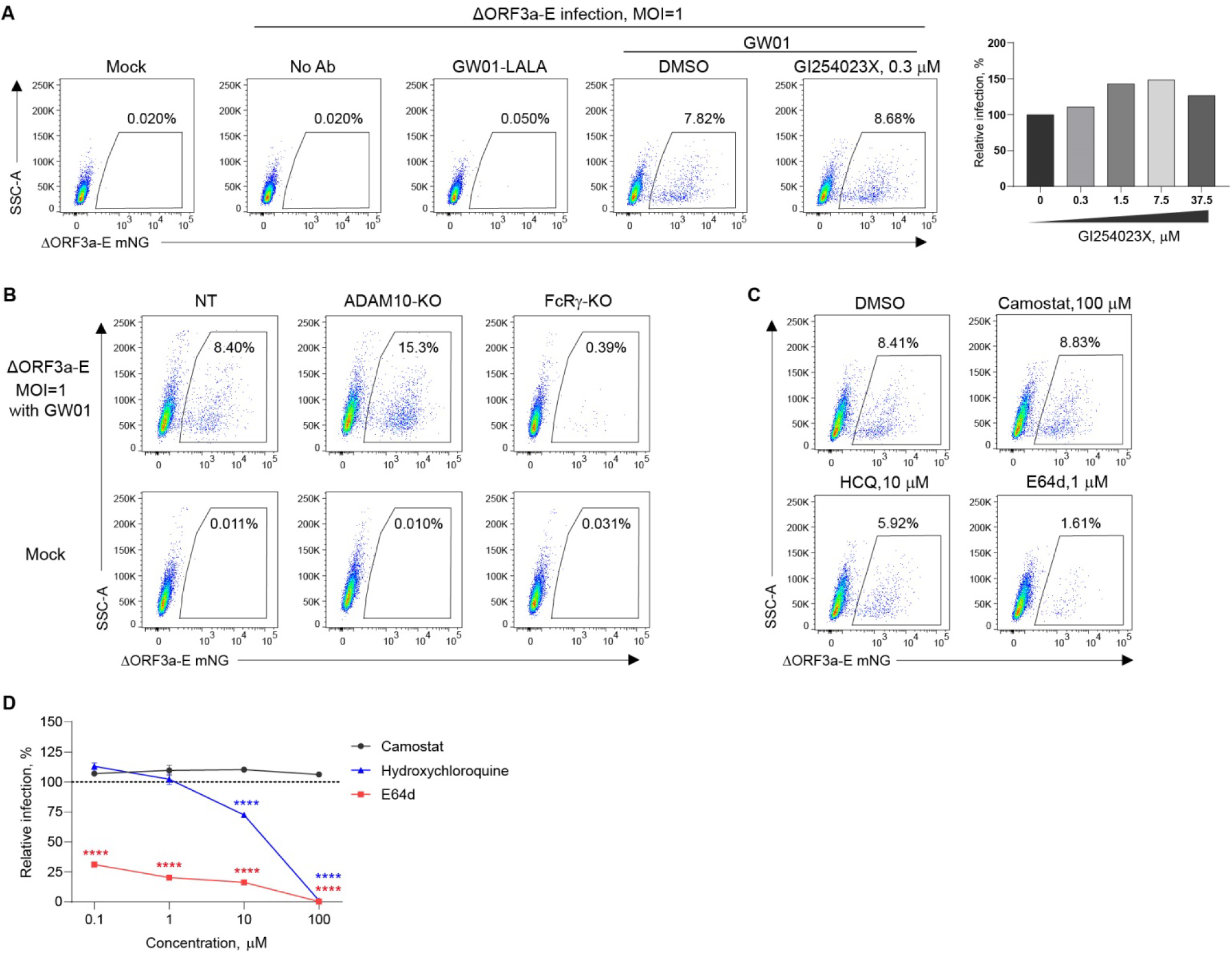
Antibody-mediated SARS-CoV-2 infection of monocytes occurs via endocytosis and is independent of ADAM10. (**A**) Representative flow cytometry plots showing the effect of GI254023X (ADAM10 inhibitor) on GW01-dependent infection of THP-1. Histogram (right) shows the effect of GI254023X concentration on SARS-CoV-2 infection of THP-1. (**B**) Representative flow cytometry dot plots showing the effect of ADAM10 or FcRγ knockout in THP-1 cells on GW01-dependent direct infection of THP-1. (**C**) Representative flow cytometry plots showing the effect of the indicated doses of camostat, hydroxychloroquine (HCQ), and E64d on GW01-dependent infection of THP-1. (**D**) Dose-dependent effect of camostat, hydroxychloroquine (HCQ), and E64d on GW01-dependent SARS-CoV-2 infection of THP-1 (mean ± SEM, n=3, ****P<0.0001, compared with camostat group at the same dose, by two-way ANOVA).

SARS-CoV-2 enters host cells via two main routes: virus-cell membrane fusion and endocytosis (23). Since antibody-mediated infection of monocytes did not depend on ADAM10 at the plasma membrane, we hypothesized that viral entry occurs via the endocytic pathway. We therefore tested endocytosis-pathway inhibitors —hydroxychloroquine, which inhibits endosomal acidification, and E64d, a cathepsin L inhibitor — on antibody-mediated infection of monocytes. For comparison, we also tested camostat, a TMPRSS2 inhibitor that blocks the cell membrane fusion pathway. Endocytosis inhibitors significantly reduced viral entry, with E64d showing the strongest effect, reducing infection by 70% at concentrations as low as 0.1 µM (Figure 5, C and D). In contrast, camostat had no inhibitory effect at any concentration tested (Figure 5, C and D). These results demonstrate that SARS-CoV-2 infection of monocytes occurs primarily through antibody-mediated endocytosis, a mechanism distinct from antibody-mediated syncytium formation.

### NAb-mediated fusion, rather than direct viral infection, is the major source of inflammation

Because antibodies mediated both syncytium formation and direct viral infection, we asked which process drives the production of these inflammatory mediators. Notably, the ADAM10 inhibitor GI254023X (blocking cell-cell fusion) potently and selectively suppressed IL-6, IL-1β, and TNF secretion in the 24-h co-culture, whereas the cathepsin L inhibitor E64d (blocking direct infection) had only marginal effects (Figure 6A). The mRNA levels of IL-6 and IL-1β were concordant with their protein levels, whereas those of TNF were not (Figure 6B), indicating that TNF is regulated at post-transcriptional stage(s). These findings demonstrate that spike antibody-induced syncytia are the major contributors to the inflammatory responses during SARS-CoV-2 infection.

**Figure 6.**
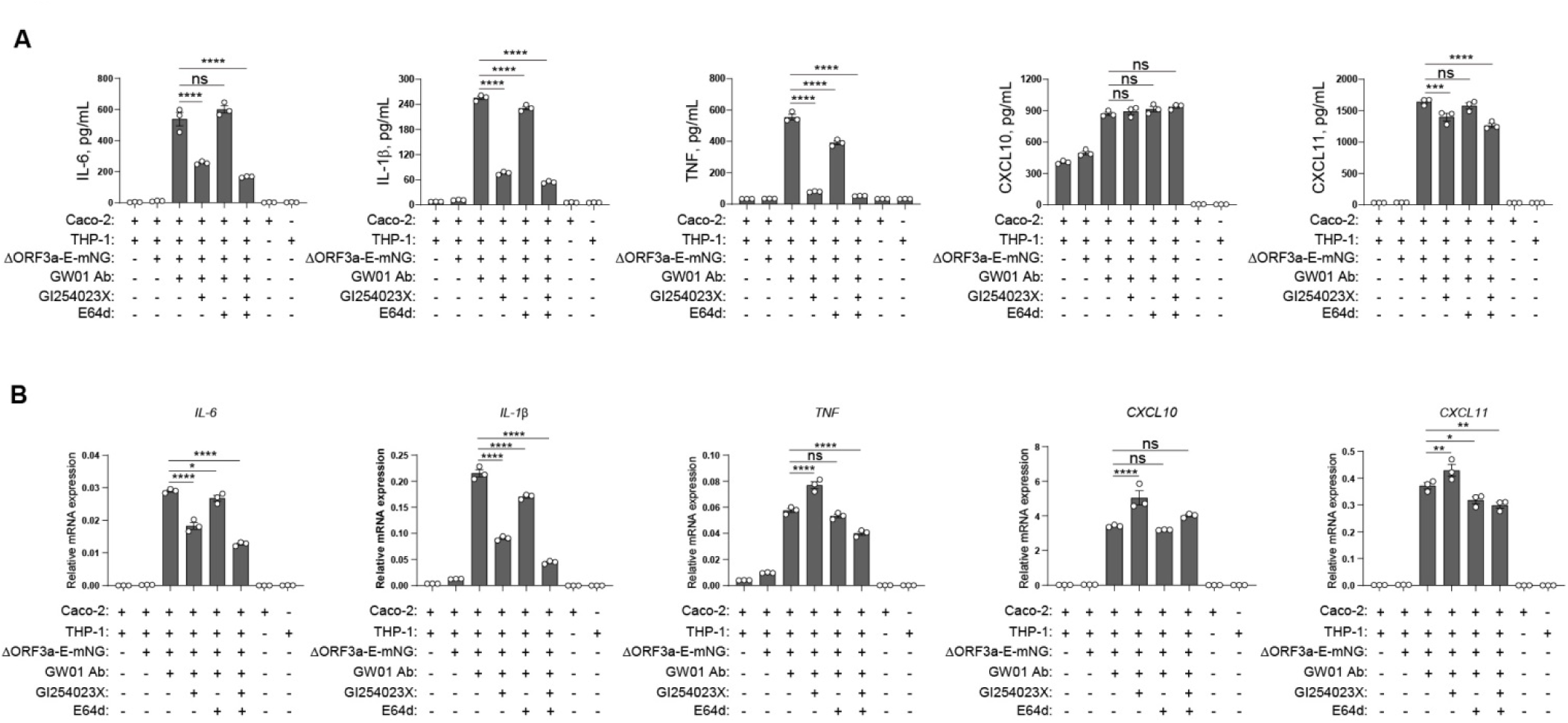
Antibody-induced fusion is the major driver of inflammation. (**A**) Effects of GI254023X and E64d on cytokine production in 24-h co-cultures measured by ELISA. This co-culture assay was modified from the method described in Fig. 3A, with the addition of inhibitors as indicated. (**B**) Cytokine mRNA quantification of co-culture from (A). Data are presented as mean ± SEM, n=3, ns means nonsignificant, *P<0.05, **P<0.01, ***P<0.001, ****P<0.0001, by one-way ANOVA.

### Hyperinflammatory SARS-CoV-2 RNA–positive syncytia are present in COVID-19 lungs

SARS-CoV-2 can trigger syncytium formation. Autopsy analysis of lungs from patients with COVID-19 reported multinucleated giant cells in ∼90% of samples (3). These multinucleated cells expressed pneumocyte markers, but whether monocytes or macrophages also participate in their formation has remained unclear. To explore *in vivo* evidence of monocyte/macrophage involvement, we reanalyzed two public scRNA-seq datasets from distinct studies: one derived from postmortem lung tissue (7), and the other from peripheral blood mononuclear cells (PBMCs), bronchoalveolar lavage fluid (BALF), and sputum (6), of COVID-19 patients. We identified a distinct doublet-like population in both datasets (Figure 7A, cluster 9, and Supplementary Figure 4A, the purple cluster) that co-expressed monocyte/macrophage marker CD163 and endothelial cell markers (Figure 7, C and D, and Supplementary Figure 4, C and D). For convenience, we refer to this population as “doublets.” Notably, SARS-CoV-2 RNA, *FCER1G* and *ADAM10* were also enriched in this population (Figure 7, B and D, and Supplementary Figure 4, B and D). These findings suggest that monocyte/macrophage-derived heterotypic syncytia may be present in the lungs of COVID-19 patients.

**Figure 7.**
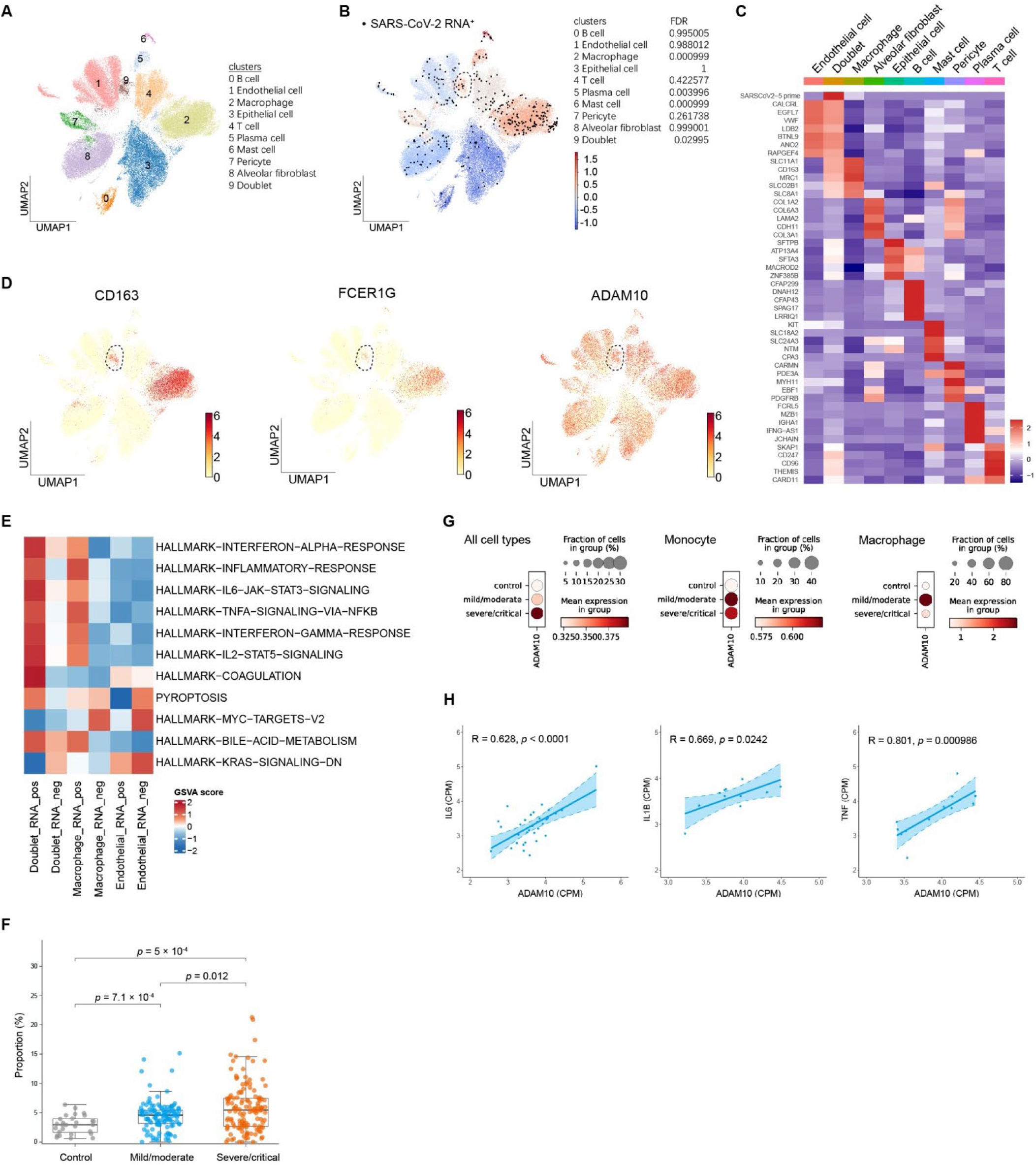
Monocyte/macrophage-derived syncytia are present in COVID-19 lungs and display a hyperinflammatory profile. (**A**) scRNA-seq clustering of postmortem lung tissue from patients with COVID-19. (**B**) SARS-CoV-2 RNA expression across clusters. (**C**) Heatmap of SARS-CoV-2 gene expression and cluster marker genes for each population. (**D**) Expression of CD163, FCER1G, and ADAM10. (**E**) Heatmap of pathway enrichment in each cell population. (**F**) Proportion of doublet in samples from severe/critical (n=134), mild/moderate (n=122), and healthy controls (n=28). (**G**) ADAM10 expression in indicated cell types across clinical severity groups. (**H**) Correlation of ADAM10 with IL6, IL1B, and TNF expression in doublets from postmortem COVID-19 lung tissue.

Pathway enrichment analysis showed that SARS-CoV-2 RNA-positive doublets exhibited significant enrichment of inflammation-related pathways, including inflammatory response, IL-6, and TNFα signaling (Figure 7E), mirroring our co-culture assay data (double-positive syncytia in the GW01-treated group; Figure 3B). SARS-CoV-2 RNA-positive macrophages also displayed moderate enrichment of inflammatory pathways (Figure 7E), consistent with GW01-treated THP-1 cells (Figure 3B). By contrast, SARS-CoV-2 RNA-negative doublets, RNA-negative macrophages, and all endothelial cells showed much lower enrichment (Figure 7E). Interestingly, coagulation pathways were also enriched in SARS-CoV-2 RNA-positive doublets (Figure 7E), with increased expression of von Willebrand factor (VWF), tissue factor (TF), and factor VIII (F8) (Supplementary Figure 4E), potentially contributing to COVID-19-associated microthrombosis (29). Notably, doublets were significantly more abundant in samples from patients with severe COVID-19 than in those from patients with mild disease or healthy donors (Figure 7F). Collectively, these observations phenocopy our co-culture results and support the conclusion that monocyte/macrophage-derived syncytia constitute a major source of inflammatory cytokines during SARS-CoV-2 infection.

We further observed that *ADAM10* expression in lung cells correlated with COVID-19 severity, with severe cases showing the highest levels across all cell types (Figure 7G and Supplementary Figure 4F). *ADAM10* was particularly enriched in monocytes and macrophages in the lungs of all COVID-19 patients, suggesting a prevalent propensity for antibody-mediated heterotypic syncytia and hyperinflammation (Figure 7G and Supplementary Figure 4F). Furthermore, expression of *IL-6, IL-1β,* and *TNF* in doublets positively correlated with *ADAM10* levels (Figure 7H). Thus, ADAM10 appears to facilitate syncytium formation between monocyte/macrophage and virus-infected pneumocyte and thereby drive pro-inflammatory responses in humans.

## DISCUSSION

Monocytes/macrophages are a significant source of pro-inflammatory cytokines in COVID-19 and are strongly linked to disease severity and mortality. Our study reveals that RBD-targeting SARS-CoV-2 neutralizing antibodies can induce fusion between virus-infected pneumocytes and monocytes/macrophages. This process begins with antibody-mediated bridging, in which the antibody simultaneously binds spike on pneumocytes and CD64 on monocytes/macrophages. ADAM10 then activates the fusion machinery, facilitating cell–cell fusion. The resulting syncytia produce large amounts of inflammatory cytokines and chemokines, which may exacerbate disease progression. Importantly, sub-neutralizing concentrations of antibody amplified inflammation and worsened disease outcomes in infected mice. Collectively, our findings identify antibody-mediated monocytes/macrophages-derived syncytia as a key mechanism driving hyperinflammation in severe COVID-19 (Supplementary Figure 5, Graphical abstract).

SARS-CoV-2 can enter cells through TMPRSS2-dependent plasma-membrane fusion or cathepsin-dependent endocytosis (23). Although monocytes/macrophages lack ACE2/TMPRSS2, scRNA-seq of COVID-19 lungs shows viral RNA enrichment in these populations (6, 7). Our data reconcile these observations by delineating two FcR-dependent routes into monocytes/macrophages: (i) cell–cell fusion, which requires CD64 and ADAM10 and generates hyperinflammatory syncytia; and (ii) endocytosis of opsonized virus, which requires CD64 and cathepsin L but not ADAM10. The former pathway links antibody specificity to tissue inflammation through a fusion mechanism; the latter provides an entry route that is comparatively less pro-inflammatory in our assays.

Our work identifies a non-canonical role for spike antibodies: promoting viral invasion of monocytes/macrophages through cell-cell fusion. This phenomenon represents a form of antibody-dependent enhancement (ADE) (30), more specifically antibody-dependent enhancement of inflammation (ADEI) (31), as it leads to massive inflammatory cytokine production by monocyte/macrophage-derived syncytia rather than increased viral replication. Unlike classical ADE in dengue, which is mediated by non-neutralizing or low-affinity neutralizing antibodies (32, 33) and enhances infection in natural target cells with robust progeny virus production (34, 35), spike antibody-mediated ADEI drives infection of non-susceptible monocytes/macrophages and amplifies inflammatory cytokines. Whether SARS-CoV-2 can productively replicate in monocytes/macrophages remains debated (12, 13). Although we did not dissect downstream cytokine mechanisms here, prior studies implicate TLR7/8, NLRP3, and AIM2 inflammasomes (12, 14, 36). Cytosolic RNA sensors such as RIG-I and MDA5 may also contribute. While most of our mechanistic work relied on monoclonal antibody assay *in vitro*, *in vivo* infection of FcγRI-humanized mice suggested that defined doses of spike antibody can enhance inflammation and worsen disease, in a polyclonal setting.

These results help explain paradoxical early-pandemic observations: severe COVID-19 patients often displayed higher neutralizing antibody titers than mild or asymptomatic cases (8, 9). We propose that infection-induced antibodies can worsen disease by promoting fusion of infected cells with monocytes/macrophages, fueling a cytokine storm. Not all antibodies possess this fusogenic property. The RBD-targeting antibodies in our study largely belong to class 3 neutralizing antibodies that bind non-RBM regions, which may favor conformational changes that promote fusion. Structural studies are needed to clarify this potential mechanism. Importantly, antibodies from natural infection often target narrower epitope repertoires (37), potentially heightening pathogenic risk if fusion-promoting clones are present. By contrast, vaccines induce broader polyclonal antibody responses (37), which may competitively inhibit fusogenic antibodies and reduce risk. Whether differences in epitope specificity between infection- and vaccine-induced repertoires contribute to divergent clinical outcomes remains to be determined.

Our findings also inform therapeutic strategies. Current antivirals such as remdesivir, paxlovid, and molnupiravir reduce viral replication and are effective in early disease (38), but they offer limited benefit in late-stage disease dominated by hyperinflammation. In such cases, immunomodulators (e.g., corticosteroids, IL-6 receptor antagonists, JAK-STAT inhibitors) are needed (38), though broad immunosuppression risks delayed clearance and secondary infections (39). Notably, our transcriptomic analyses of COVID-19 patient samples revealed that monocyte/macrophage-derived syncytia exhibit a pronounced pro-inflammatory signature, whereas infected epithelial/endothelial cells alone showed little enrichment of inflammatory pathways. Pharmacological inhibition of monocyte/macrophage fusion with an ADAM10 antagonist reduced secretion of IL-6, IL-1β, and TNFα *in vitro*. We therefore propose that ADAM10-dependent monocyte/macrophage fusion is a central contributor to excessive inflammation in severe disease and that directly inhibiting monocyte/macrophage-derived syncytium formation could mitigate cytokine storms at their source. Prior work showed ADAM10 blockade prevents epithelial syncytium formation (40). Thus, ADAM10 inhibitors may offer a dual benefit by limiting viral spread and dampening hyperinflammation without global immunosuppression. ADAM10 is a broadly active metalloproteinase with diverse substrates, such as Notch1/2, multiple adhesion molecules, and amyloid precursor protein, thereby may contribute to the regulation of immune responses, tumorigenesis, and Alzheimer’s disease (28). It is normally autoinhibited at the plasma membrane (41), and its activation is regulated by multiple events such as intracellular domain phosphorylation or adaptor protein interactions (42, 43), the details of which remain unclear. Thus, the precise mechanism of ADAM10 activation during antibody-mediated monocyte/macrophage fusion remains an open question warranting further study.

In summary, we demonstrate that certain spike RBD-specific monoclonal antibodies drive syncytium formation between SARS-CoV-2-infected epithelial cells and monocyte/macrophage through an FcγRI (CD64) and ADAM10-dependent mechanism. These syncytia produce excessive inflammatory cytokines and chemokines, and antibodies at suboptimal doses can exacerbate disease in vivo. This work identifies antibody-mediated heterotypic cell fusion as a previously unrecognized contributor to COVID-19 severity and nominates ADAM10 as a potential therapeutic target for severe disease.

## METHODS

### SARS-CoV-2 infection and GW01 treatment in FcγRI-humanized mice

C57BL/6NCya-Fcgr1tm1(hFCGR1A)/Cya (hFCGR1; Cyagen, C001500) mice were housed at the Louisiana State University School of Veterinary Medicine. All procedures were approved by the IACUC. Twelve-week-old mice were challenged intranasally with 10^5 PFU of mouse-adapted SARS-CoV-2 (MA10; BEI Resources, NR-55329) in 50 µL sterile PBS, followed 2 days later by retro-orbital intravenous administration of GW01 (dose as indicated) in 100 µL sterile vehicle. Animals were monitored daily. At 6 days post-infection (dpi), mice were euthanized and lungs were collected. Portions were fixed in 10% neutral-buffered formalin for ≥72 h, processed, and embedded for H&E staining. Pathology scoring was performed by a board-certified veterinary pathologist as previously described (44).

### Cell lines

A549, THP-1 and HEK293T cells were obtained from ATCC. Caco2-3a-E cells are Caco-2 stably expressing SARS-CoV-2 ORF3a and E and an mCherry reporter under a doxycycline-inducible promoter (21)). A549 cell line stably expressing SARS-CoV-2 spike (A549-spike) was generated by lentiviral transduction and blasticidin selection. A549-spike, HEK293T, and Caco2-3a-E cells were maintained in complete DMEM (10% heat-inactivated FBS), 1% penicillin and streptomycin (P/S)). THP-1 cells were maintained in complete RPMI-1640 (10% FBS, 1% P/S).

### Antibody-mediated cell-cell fusion/syncytium formation assay

A549-spike cells were plated in 6-well plates at 0.3 × 10^6 cells per well in complete DMEM. After overnight incubation, cells were washed twice with PBS and treated with spike-neutralizing antibody GW01 (or indicated clones) at 1 µg/mL; untreated cells served as controls. After 30 min, 1 × 10^6 THP-1 cells were added per well for co-culture. At 48 h, cultures were examined and imaged by bright-field microscopy. For flow cytometric quantification, A549-spike cells were labeled with CFSE (5 µM; eBioscience, 65-0850-84) before plating. The next day, THP-1 cells were labeled with eFluor450 (5 µM; eBioscience, 65-0842-90) and added to A549-spike cells. After 24 h, adherent cells were trypsinized, combined with supernatants, washed once with staining buffer (PBS + 2% FBS + 1 mM EDTA), resuspended, and acquired on a BD LSRFortessa. Data were analyzed in FlowJo v10.8.1.

### Live cell imaging of antibody-induced fusion

A549-spike cells were labeled with CFSE (5 µM), seeded in 24-well plates (0.06 × 10^6 cells/well), and cultured overnight. After two PBS washes, cells were treated with GW01 (1 µg/mL) for 30 min. THP-1-mCherry cells (0.2 × 10^6 per well; pretreated with GI254023X where indicated) were added, and plates were immediately transferred to an Incucyte® SX1 (Sartorius) for real-time imaging with a 20× objective.

### Primary human monocytes and monocyte-derived macrophages (MDMs)

Peripheral blood from healthy donors was obtained from the Shanghai Blood Center. Human peripheral blood mononuclear cells (PBMCs) were obtained by density gradient centrifugation with Lymphoprep (Stemcell, 07801). Primary monocytes were isolated form PBMCs using negative selection (Stemcell, 19359). For MDMs, monocytes were cultured at 0.5 × 10^6 cells/mL in complete RPMI-1640 with either GM-CSF (25 ng/mL; PeproTech, 300-03-20UG) or M-CSF (50 ng/mL; Gibco, PHC9501) for 7 days at 37 °C, 5% CO2; medium was replaced every 2–3 days with fresh cytokine-containing medium. Monocytes, GM-CSF–MDMs (G-MDMs), and M-CSF–MDMs (M-MDMs) were assayed for antibody-mediated syncytium formation and quantified by flow cytometry as above.

### NanoLuc-based cell-cell fusion assay

A549-spike-HiBiT cells were resuspended in complete culture medium and seeded in 96-well plates at 0.01 × 10^6 cells per well. After overnight culture, the culture medium was aspirated, and 50 µL of spike-specific antibody clone GW01 (4 µg/mL in culture medium) was added to each well. Following 30 min of incubation, 50 µL of THP-1-LgBiT cells (0.02 × 10^6 cells/well) were added, and co-cultured overnight. The next day, 25 µL of Nano-Glo® Live Cell Reagent (Promega, N2012) was added, and luminescence (RLU) was measured using a Synergy H1 microplate reader. The assay was performed according to the manufacturer’s instructions.

For antibody titrations, A549-spike-HiBiT cells were treated with serial dilutions (final: 0.000128, 0.00064, 0.0032, 0.016, 0.08, 0.4, 2, 10, 50 µg/mL), followed by THP-1-LgBiT addition and luminescence readout as above.

For EK1C4 inhibition, A549-spike-HiBiT cells were pretreated with EK1C4 (1, 10, 100 nM) for 30 min, then incubated with serial GW01 (0.000128–50 µg/mL) and co-cultured overnight with THP-1-LgBiT before luminescence measurement.

To assess ADAM10 and FcγR requirements, THP-1-LgBiT cells were pre-incubated for 30 min with GI254023X (9.375–9,600 nM) or CD64/CD32 blocking antibodies (3.906–4,000 ng/mL), then co-cultured with A549-spike-HiBiT. Luminescence was quantified as above.

### Genome-wide CRISPR screening for fusogens

The genome-wide CRISPR sgRNA library (Addgene, 101926-101934) was packaged into lentiviruses. THP-1-Cas9 cells were transduced at an MOI of 0.3 and selected with puromycin (1 µg/mL, 5 days) to generate THP-1-lib cells (stable sgRNA library; mCherry^+^). A549-spike cells were pre-treated with GW01 (1 µg/mL, 30 min) and co-cultured with THP-1-lib for 24 h (Round 1). Unfused THP-1-lib cells in the supernatants were collected and re-co-cultured with fresh GW01-treated A549-spike cells for two additional 24-h rounds. In Round 3, A549-spike cells were pre-labeled with CFSE. After three rounds, pooled cells (supernatant + adherent) were sorted on a BD FACSAria III into mCherry^+^ CFSE^-^ (unfused THP-1-lib) and mCherry^+^ CFSE^+^ (syncytia). Genomic DNA was extracted (Qiagen, 51192); sgRNA regions were PCR-amplified with NEB HiFi Master Mix (M0543L), gel-purified (2% agarose), and sequenced on an Illumina NovaSeq 6000 (∼60 million reads/sample). sgRNA enrichment in unfused vs syncytia gates was quantified with MAGeCK; hits were ranked by fold-change and *p*-value.

### Functional validation of CRISPR-screened candidate genes

sgRNAs targeting ADAM10 and FCER1G were cloned into pMCB320 (Addgene, 89359) and used to transduce THP-1-Cas9 cells (third-generation packaging). After puromycin selection (1 µg/mL, 7 days) and cloning, three independent knockouts per gene were established (ADAM10-KO-1/-2/-3; FcRγ-KO-1/-2/-3). Non-targeting (NT) sgRNA cells served as controls. Antibody-dependent fusion assays were performed as above.

For rescue, codon-optimized ADAM10 and FCER1G cDNAs (with synonymous mutations at sgRNA sites) were cloned into pLVX-IRES-hygro, packaged into lentivirus, and used to complement knockouts (ADAM10 cDNA into ADAM10-KO; FcRγ-cDNA into FcRγ-KO). Empty-vector controls (NT-LVX, ADAM10-KO-LVX, FcRγ-KO-LVX) were included. Fusion assays were repeated to confirm rescue.

### ORF3a-E mNG virus production and titration

Caco-2-3a-E cells were seeded in T175 flasks and induced with doxycycline (200 ng/mL). After 24 h, medium was replaced and cells were infected with ΔORF3a-E-mNG at an MOI of 0.01. Viral supernatants were harvested 72 h post-infection. For titration, Caco-2-3a-E cells were plated in 96-well plates (0.02 × 10^6 cells/well) and induced with doxycycline (200 ng/mL) overnight. Serial virus dilutions (100 µL/well; n = 6 per dilution) were applied; after 48 h, mNG-positive foci were counted by fluorescence microscopy to calculate the viral titer.

### Antibody-mediated SARS-CoV-2 infection of THP-1

Unless otherwise specified, THP-1 cells were infected at an MOI of 1. ΔORF3a-E-mNG was pre-incubated with GW01 or GW01-LALA (0.2 µg/mL) for 30 min (37 °C, 5% CO2) and mixed with 0.1 × 10^6 THP-1 cells. At 24 h post-infection, mNG^+^ cells were quantified by flow cytometry.

For ADAM10 inhibition, THP-1 cells were pretreated with GI254023X for 30 min before infection. To test entry inhibitors, THP-1 cells were pre-incubated with camostat, E64d, or hydroxychloroquine (0.1, 1, 10, 100 µM) for 30 min before exposure. For genetic tests, ADAM10-KO or FcRγ-KO THP-1 (0.1 × 10^6) were infected with GW01-pretreated ΔORF3a/E-mNG (0.2 µg/mL, 30 min) and analyzed at 24 h.

### Spike antibody-induced syncytia during infection, cell sorting, and bulk RNA-seq

Caco-2-3a-E cells were seeded in 6-well plates (0.2 × 10^6 cells/well) in medium with doxycycline (200 ng/mL) overnight, infected with ΔORF3a-E-mNG (MOI=1) for 1 h, after which medium was replaced. The next day, cells were treated with GW01 (1 µg/mL, 30 min) and co-cultured with THP-1 cells (1 × 10^6 per well). After 24 h, supernatants and cells were collected for cytokine and RNA analyses.

To assess inhibitors, THP-1 cells were pre-incubated with GI254023X (10 µM), E64d (1 µM), or both for 30 min before co-culture with infected, GW01-treated Caco-2-3a-E cells.

For population-specific RNA-seq, Caco-2-3a-E cells were labeled with eFluor450 (5 µM) before seeding, infected (MOI=1), treated with GW01 (1 µg/mL, 30 min), and co-cultured with THP-1 for 24 h. Cells were stained with anti-CD64-APC and sorted on a FACSAria III into: syncytia (eFluor450^+^ APC^+^), unfused Caco-2-3a-E (eFluor450^+^ APC^-^), and unfused THP-1 (eFluor450^-^ APC^+^). Total RNA was extracted (Qiagen, 74134) and libraries prepared (Illumina, 20020189) according to the manufacturer’s instructions. Libraries were sequenced on an Illumina NovaSeq X Plus (paired-end 150 cycles). Adaptor/low-quality reads were removed with Trimmomatic v0.36 (45). Clean reads were aligned to Ensembl GRCh38 with default parameters in STAR v2.7.0 (46). Gene counts were generated with featureCounts v2.0.1 (47). Pathway enrichment was performed with the GSVA function (GSVA R package v1.38.2) (48).

### Single cell RNA-seq data analysis

Raw sc/snRNA-seq data were retrieved from DUOS and aligned with Cell Ranger (10x Genomics) to custom references (GRCh38_and_SARSCoV2 for scRNA-seq; GRCh38_premrna_and_SARSCoV2 for snRNA-seq). The pre-mRNA reference captures intronic and exonic reads. The SARS-CoV-2 FASTA/GTF were as previously described; the GTF was edited to include CDS only plus 5′ UTR (“SARSCoV2_5prime”), 3′ UTR (“SARSCoV2_3prime”), and negative-strand regions (“SARSCoV2_NegStrand”). Cells with <100 genes, <500 UMIs, or >15% mitochondrial reads were removed. To mitigate batch effects, Seurat FindIntegrationAnchors (default) was applied across patients; likely multiplets were identified and removed. Seurat v4.0 (49) was used for normalization, clustering, and differential expression (defaults unless stated). For all-cell clustering, 2,000 variable genes were selected, PCA was performed (RunPCA), and the top 20 PCs were used for clustering. DEGs were identified with FindAllMarkers/FindMarkers (adjusted P < 0.05). Viral enrichment was performed as described (7). Pathway analyses used GSVA (v1.38.2). Spearman correlations were used to relate ADAM10 to cytokine genes.

### RT-qPCR

Total RNA from cells was extracted with Direct-zol RNA Miniprep (Zymo, R2051), reverse-transcribed with PrimeScript RT (TaKaRa, RR047A), and analyzed by qPCR using TB Green Premix Ex Taq II (TaKaRa, RR820A). Gapdh was used as the housekeeping control, and relative expression was calculated by the 2^–ΔCt method.

For lung tissue, the post-caval lobe was weighed and homogenized in PBS (1 mL, pH 7.4) using a TissueLyser LT (Qiagen). RNA was extracted with TRIzol™ LS (Invitrogen) and cDNA synthesized with ProtoScript II (NEB). SARS-CoV-2 genomes were quantified using the 2019-nCoV RUO Kit (IDT, 10006713) with a standard curve generated from Heat-Inactivated SARS-Related Coronavirus 2, USA-WA1/2020 (BEI Resources, NR-52347). Cytokine transcripts were quantified with TaqMan “Best Coverage” assays for mouse Gapdh (Mm99999915_g1), Il6 (Mm00446190_m1), Tnf (Mm00443258_m1), Cxcl11 (Mm00444662_m1), and Cxcl12 (Mm00445553_m1). Values were normalized to Gapdh and to the mean of uninfected controls.

### Cytokine ELISA

Human IL-6, IL-1β, TNF-α, CXCL10, and CXCL11 in culture supernatants were quantified with commercial ELISA kits according to the manufacturer’s protocols: IL-6 (Dakewe, 1110602), IL-1β (Dakewe, 1110122), TNF-α (Dakewe, 1117202), CXCL10 (Neobioscience, EHC157.96), CXCL11 (Neobioscience, EHC084.96).

### Statistics

Statistical analyses were performed using GraphPad Prism 8.0.2 software. Comparisons between two groups were made using Student’s t-test, while comparisons among multiple groups were evaluated using one-way or two-way ANOVA, as indicated in figure legends. Data are reported as mean ± SEM, \**P* < 0.05, \*\**P* < 0.01, \*\*\**P* < 0.001, and \*\*\*\**P* < 0.0001.

## Data availability

Raw human sequencing data were retrieved from the controlled access repository Data Use Oversight System (https://duos.broadinstitute.org/, IDs: DUOS-000126, DUOS-000127, DUOS-000128 DUOS-000129) and Genome Sequence Archive (https://ngdc.cncb.ac.cn/gsa-human, ID:HRA001149).

Viral genome assemblies and short-read sequencing data are available in NCBI’s GenBank and SRA databases, respectively, under BioProject PRJNA720544. GenBank accessions for SARS-CoV-2 genomes are MW885875–MW885883. Processed bulk RNA-seq profiles generated during this study are publicly available at https://zenodo.org/records/17139396.

## Author contributions

YY, CZ, TZ, WH, NM, MC, NJ, TL, YW, and CW conducted the experiments. YY, CZ, and TZ analyzed data. FW, WH, JH, YY, CZ, and TZ designed the experiments. FW, WH, and JH supervised the conduct of the study. FW and YY wrote the initial draft of the manuscript. FW and WH revised the manuscript. YY, CZ, and TZ made significant contributions to this work and share first authorship.

## Funding Support

National Key Research and Development Program of China (2023YFE0118200 to F.W., 2025YFC2311600 to J.H.).

National Youth Talent Support Program of China (F.W.).

“Shuguang Program” of Shanghai Education Development Foundation and Shanghai Municipal Education Commission (22SG16 to F.W.).

Fundamental & Clinical Collaborative Innovation Team Program of Shanghai Cancer Institute (SMC-BCC-02 to F.W.). 2024 Hevolution/AFAR New Investigator Award (HEV∼NI24020 to W.H.). National Natural Science Foundation of China (32370995 and 31771008 to J.H.).

Shanghai Municipal Science and Technology Major Project (ZD2021CY001 to J.H.). Zhejiang Provincial Natural Science Foundation of China (LY23H100001 to Y.Y.).

National Key Research and Development Program of China (2023YFC2509600 to C.Z.). Postdoctoral Innovation Talent Support Program (BX20240084 to Y.W.).

## Acknowledgments

We thank the staff of the Core Facility of Shanghai Immune Therapy Institute for cell sorting and imaging assistance. We also thank Dr. Qiming Liang and Xin Wang from Shanghai Institute of Immunology for providing the ΔORF3a-E mNG virus and Caco2-3a-E cell line, as well as technical guidance on the viral amplification; Dr. Lu Lu from Fudan University for providing the EK1C4 peptide; and Dr. Avery August at Cornell University for critical comments and feedback.

