## Supplemental data for "Antibody-Dependent Heterotypic Syncytia Drive COVID-19 Inflammation and Disease Progression"

**The PDF file includes:**

Supplementary Figure 1 to 5

**Other Supplementary Materials for this manuscript include the following:**

Movies S1 to S3

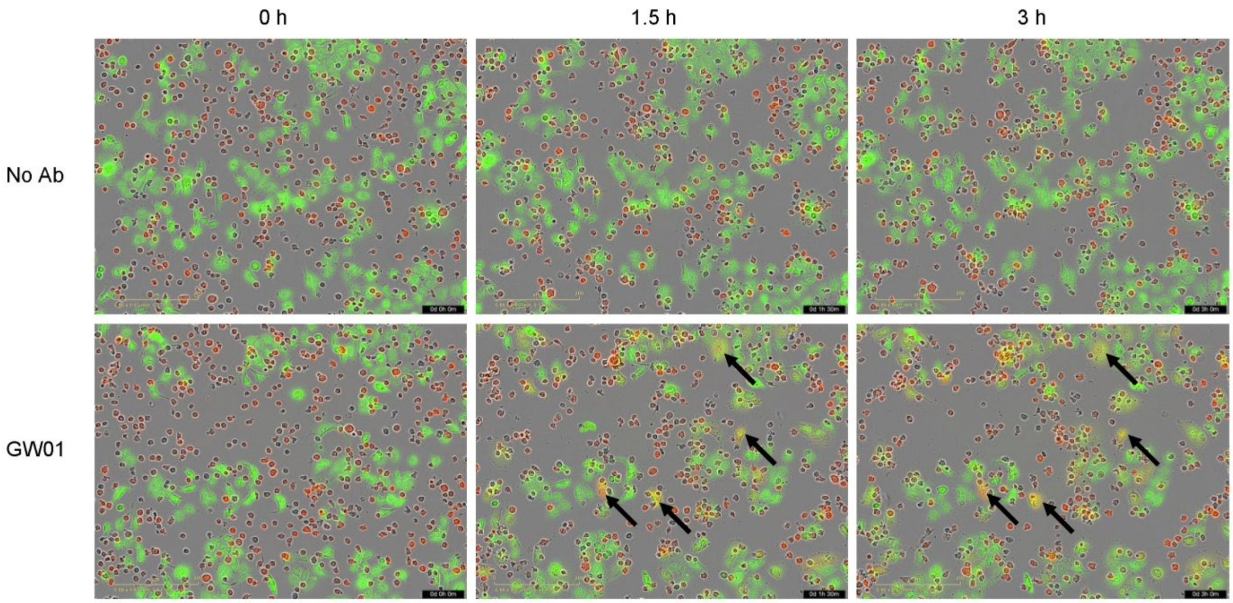


**Supplementary Figure 1**

Spike antibody GW01 induces syncytium formation. A549-spike (green) and THP-1(red) cells were co-cultured, with or without GW01 antibody, and syncytium formation was recorded with an Incucyte imaging system. Black arrows indicate syncytia.

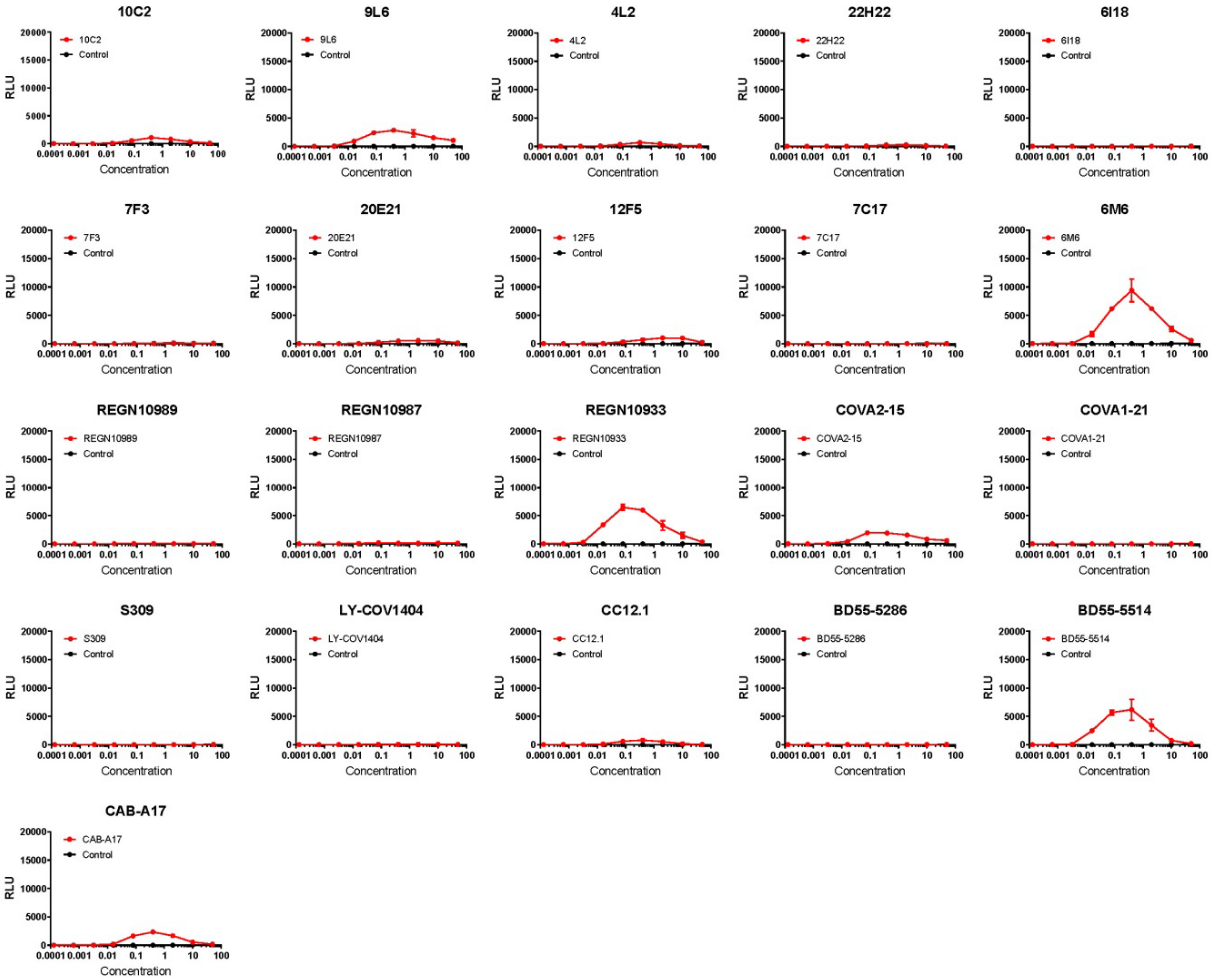


**Supplementary Figure 2**

Syncytium-forming activity of spike monoclonal antibody clones. Individual spike monoclonal antibodies were evaluated with a Nanoluc assay alongside a control antibody to test their ability to induce syncytia between THP-1 and A549-spike cells.

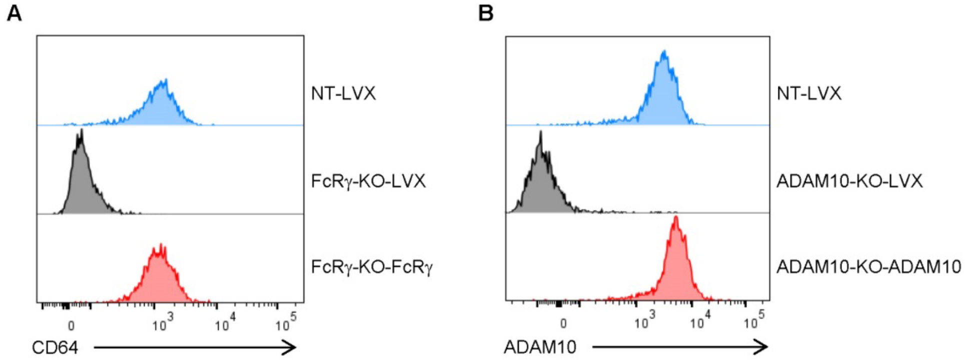


**Supplementary Figure 3**

Deletion and restoration of CD64 and ADAM10 in THP-1. FACS histograms showing the expression of CD64 (A) and ADAM10 (B) in the indicated cells.

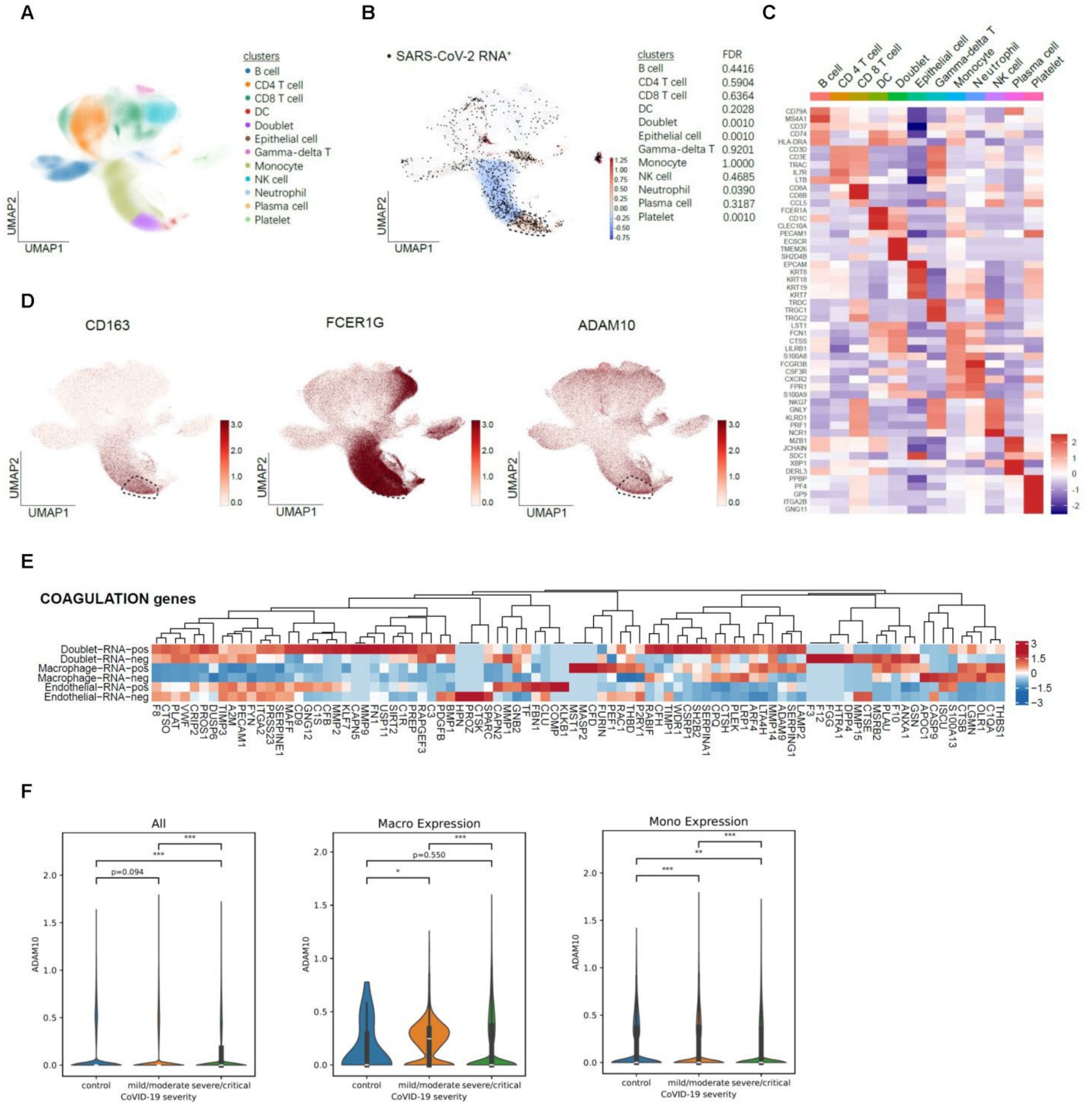


### Supplementary Figure 4

Monocyte/macrophage-derived syncytia in COVID-19 samples, the association between ADAM10 expression and COVID-19 severity, and expression of coagulation pathway genes in the indicated cell clusters. (A) scRNA-seq clustering of cells derived from PBMC, BALF, and sputum samples from patients with COVID-19 and healthy donors (n=284, total). (B) Expression of SARS-CoV-2 RNA across cell clusters. (C) Heatmap showing marker-gene expression in each cluster. PECAM1 and ECSCR are markers of endothelial cells. (D) Expression of CD163, FCER1G, and ADAM10. (E) Expression of coagulation pathway genes in the indicated cell clusters with or without detectable SARS-CoV-2 RNA. (F) Violin plots showing ADAM10 expression in all cell types, macrophages, and monocytes from healthy controls and patients with mild/moderate or severe/critical COVID19.

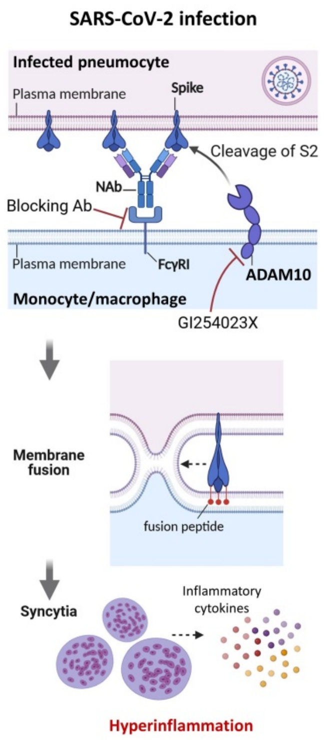


**Supplementary Figure 5**

During SARS-CoV-2 infection, certain spike antibodies mediate intercellular bridging between monocyte/macrophage and infected pneumocyte. Subsequently, the metalloproteinase ADAM10 expressed by monocyte/macrophage activates spike, which triggers cell-cell fusion and facilitates entry of viral components into monocytes/macrophages. The resulting heterotypic syncytia secrete substantial amounts of inflammatory cytokines, thereby exacerbating inflammation and disease severity. (Created with Biorender.com)

**Movie S1.**

Time-lapse video of the co-culture of THP-1-mCherry (red) and A549-spike (green, labeled with CFSE), without GW01 antibody.

**Movie S2.**

Time-lapse video of the co-culture of THP-1-mCherry (red) and A549-spike (green, labeled with CFSE), with the addition of GW01 antibody.

**Movie S3.**

Time-lapse video of the co-culture of THP-1-mCherry (red) and A549-spike (green, labeled with CFSE), with the addition of GW01 antibody and GI254023X (ADAM10 inhibitor).
